# Lecture capture – an engagement marker for student success?

**DOI:** 10.1101/2025.10.20.683432

**Authors:** Louise Robson

## Abstract

The impact of student use of lecture capture on their learning sits in a contested space, with previous publications showing evidence for improvement, no change and a decline in academic attainment by students. Given the diverse nature of our student cohorts, and the complex interplay of factors that determine success, it is not surprising that such variation exists in the literature. Therefore, rather than try to evaluate the impact of lecture capture, I wanted to determine whether use of lecture captures could be a marker for predicting student success in a second year undergraduate biomedical cohort. Total and unique lecture capture views, along with virtual learning environment (VLE) hours (a known engagement marker) were mapped across the academic year and compared to performance in assessments. The study identified that student engagement with lecture capture (total views and unique views) and VLE hours were positively correlated with exam attainment for all time points across the academic year. Students with higher grades were more likely to have higher total and unique views, while those with the lowest grades were less likely to have watched lecture captures and more likely to have binge watched captures just before the exam. However, it was also clear that some students did well even with low lecture capture views. Interestingly, when views were normalised to individual totals, it was clear that students who completed the majority of their own individual views in the teaching period (irrespective of total number) were more likely to do well in assessments. The study provides evidence that engagement with lecture capture and the VLE during teaching weeks can be a predictor of student success and aligns with the guidance given to undergraduate students around spacing their engagement with teaching resources.

## Introduction

Lecture capture, the recording of live teaching sessions so students can revisit material, has been in use for many years across the global higher education sector [1]. Its use has grown significantly over the years, with universities taking an institutional approach to the provision of lecture capture. In the UK this growth can be clearly documented through the Heads of eLearning Forum (HeLF) survey. In 2017, 86% of responding UK institutions had lecture capture, although only 38% had fully implemented it, and 19% had no formal implementation [2]. In contrast, the same HeLF survey in 2024 highlighted that 78% of responding universities had full implementation, and all universities had some implementation [3]. There have also been changes in how lecture capture is implemented institutionally, with 54% of responding institutions adopting an opt in policy and 24% opt out in 2017, versus 35% opt in and 50% opt out in 2024. This shift away from opt in to opt out clearly has the potential to lead to more sessions being recorded, and probably reflects the fact that institutions are responding to student demand for more lecture captures. The increased provision of recorded lectures has undoubtedly been influenced by the COVID-19 pandemic, where recorded lectures and bespoke recordings became the norm for how content was delivered during the online teaching period [4], leading to an increase in demand for recordings on the return to face to face teaching.

Lecture capture is almost exclusively viewed as positive by students, as reported in many studies looking at student reported use and perception. Some of these studies were focussed on small to medium or disciplinary specific cohorts [5–14], while others took an institutional approach to evaluating the student viewpoint [15, 16]. However, irrespective of the size of the cohort, disciplinary area and global location, there are some common findings across these studies:

1. Students feel that access to lecture capture helps with their learning.
2. Students value the opportunity to revisit lecture content they missed or did not understand / found difficult while in the live lecture.
3. Students value the opportunity to catch up with lectures they may have missed due to reasons such as illness, working, caring responsibilities.
4. Students state that access to lecture captures helps with revision.
5. Students state that there is minimal impact on their attendance at live lectures.

There is also a growing body of evidence that lecture capture provides critically important support for those students with additional support needs, e.g. specific learning differences, students with a disability, with caring responsibilities, students who have to work, and those where the language of instruction is not their first language [5, 15, 17, 18]. In a paper published in 2024, responses were gathered from 319 students who self-identified as neurotypical, neurodiverse or disabled, or both neurodiverse and disabled [19]. Two of themes that the study identified was that lecture capture is an inclusive tool and provides a flexible safety net. Those students who were neurodiverse and / or had a disability provided some incredibly powerful examples and quotes on the benefit of lecture capture, e.g. their disability impacting on their ability to attend, that they found it difficult to focus for a full lecture or that they often found attending sessions overwhelming.

Clearly students feel that having access to lecture capture is critically important to their learning experience. However, this overwhelmingly positive student viewpoint is in contrast to academic staff, who raise a number of concerns. The main ones reported are: lecture captures will lead to a drop in attendance; lecture captures may lead to superficial learning; recordings may impact teaching style [15, 16, 20, 21]. The literature itself is quite mixed in terms of addressing these concerns. For example, when considering attendance, two systematic reviews of the literature suggest there is minimal impact on attendance [22, 23], while some individual studies show evidence for a small drop [16, 24]. In terms of attainment, the published literature is also mixed, with different studies showing evidence for no impact of lecture capture use on attainment [25, 26], others showing a decrease in attainment [27, 28], and some showing a positive impact [29, 30]. These mixed findings are not that surprising, given that the impact of lecture capture on student attendance, learning and attainment is not something that can be simply defined, but is a complex integration of a range of factors that includes (but is not limited to) the learning and teaching environment, disciplinary area, year of study, the strategies students use in their learning and the individual student lived experience.

Interestingly, the majority of studies looking at the impact of lecture capture have tended to report student declared use of lecture captures, rather than actual use based on platform analytics. These data are therefore relying on accurate student reporting, and there may be a number of reasons why students may not accurately report on their use of lecture captures, e.g. reporting influenced by how students think their peers are using lecture captures [31]. There are a small number of papers that have aimed to directly observe student use of lecture captures, and then relate this to student achievement. In a large first year biology cohort actual views of lecture captures was significantly, positively correlated with exam performance, although the relationship was not strong [32]. In this study they also observed that those students who initially accessed the captures more than 48 hours before the mid-term exam (week 5 in a 10 week course) did better than predicted in their assessment. In contrast, those students who only accessed their first capture just before the midterm exam did worse than predicted. A year 2 study [27] documented lecture capture views at two time points; the end of teaching and the day after the exam. There were a total of 230 views by the end of the teaching period, with only 32% of the cohort using the captures. By the day after the exam the views had increased to 730, i.e. around two thirds of the views were in the revision period, from 59% of the cohort. This cohort, therefore, showed evidence of binge watching for revision purposes. Teaching period use of lecture captures in this study was positively related to coursework grade and lecture attendance, so students who used the lecture captures during the teaching period were more likely to be those who were attending. However, lecture capture views were not related to exam performance. The actual views (recorded at the end of the assessment period) of a large cohort of year 2 and 3 students in biology was also not correlated with exam performance [33]. However, in this study there was no data on the timing of how students had used the captures, with the analysis grouping those who had spaced their use across the teaching period and those who had binge watched just before the assessment into the same group. This study also grouped both year 2 and 3 students together. A study from 2024 did evaluate the “when viewed” aspect with a cohort of accountancy students [34]. They divided students into three groups: near event viewing (NEV), get round to viewing (GRV) and revision time viewing (RTV). They found that year 1 students were more likely to use the captures for revision purposes (RTV), with large increases in minutes viewed just before exams. There was no significant difference in the final year mark across the different types of lecture capture users, although there was a trend for those who had more revision period views to do less well. For year 2 students, those with a GRV approach tended to do better in the January and final year assessments. For year 3 those with a NEV approach had higher grades than the other cohorts, and there was a trend for those with RTV to have the lowest grades.

These studies show the importance of looking at actual lecture capture use, but also highlight that the timing of when the lecture captures are used and looking at year groups separately is also critically important. Some of the above studies have not considered the specific timing aspect, others have. What has not been completed is a systematic mapping of the engagement with lecture captures across a whole academic year and evaluating whether there is a correlation between the use of lecture captures and academic attainment. The study aimed to complete this mapping and analysis, and also compare engagement with lecture captures to hours spent on the virtual learning environment (VLE). Use of the VLE is a known marker for engagement, and is associated with higher academic attainment [35–38]. Evaluation of VLE hours alongside the lecture capture therefore provides a way of comparing the use of lecture capture with a known engagement marker. The overall aim of the study was to identify whether the use of lecture captures can be used as marker for engagement and a predictor of student success.

## Methods

### Module information

Data was collected from two 80 credit, academic year long modules in year 2 of a Biosciences programme (2/3^rds^ of the total student learning activity). The structure and assessment methods of these modules was the same, and so the data was collated and is presented as a single dataset. 172 students in total were registered on both modules.

A range of different assessment methods were utilised in the modules to ensure students met the learning outcomes: 1. An invigilated multiple choice examination (MCQ) common to both modules, held in the first (January) assessment period (168 students). 2. An invigilated short answer question and essay examination (SAQ/essay), held in the second assessment period (May, 165 students). This assessment tested knowledge, understanding and the ability of students to apply this to solve problems. It contained some questions common across both modules, but also questions from relevant specialist areas. A mean grade for coursework was also derived from a literature review and laboratory report (completed by 169 students). While all 172 students contributed to the whole cohort use data, for each assessment, only the analytics data for those students who completed the specific assessment was used. Students were placed into the following attainment groups: 70+, 60 to 69, 50 to 59, 40 to 49 and <40 (standard English HE grade boundaries).

### Analytics download

Lecture capture analytics were accessed from the Echo360 lecture capture platform. All lecture sessions were automatically recorded, and were available within 30 minutes after the end of the lecture. The specific analytics analysed were:

1. The number of unique lecture capture views.
2. The total number of lecture capture views.

One module had 114 lecture captures by the end of the academic year, and the second module had 113 lecture captures. All students had experience of using lecture captures in their previous year of study, and received guidance on how to use captures effectively [39].

The hours spent on the VLE (Blackboard) were also recorded for all time points. These hours did not include the time spent viewing lecture captures, as these were hosted and viewed on the Echo360 platform (outside of Blackboard). There may be a small contribution to the VLE hours as students accessed the Echo360 platform via the Blackboard page for the module.

Data was collected in weeks 4, 6, 9 and 12 in each semester. Data was also collected just before each examination. In semester 2 an additional time point was recorded at the end of the Easter vacation (which ran between weeks 9 and 10 of teaching). Table 1 shows the timing of teaching, assessment and vacations across the academic year.

**Table 1.**
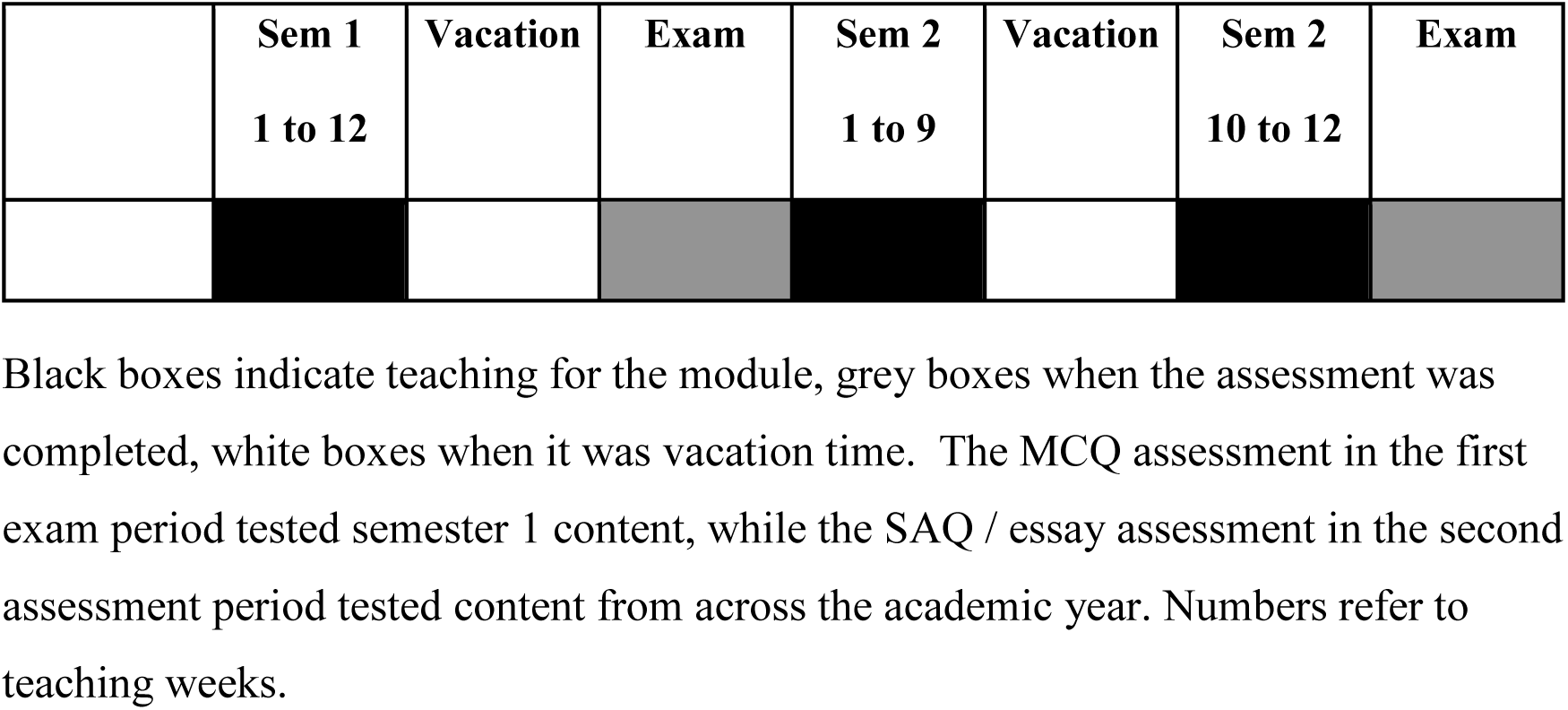
Map of teaching, examination periods and vacations for the modules.

### Statistical analysis

Data is presented as mean ± SEM. Cohort use of lecture captures (total views, percentage of unique captures viewed) across the year was evaluated using the Kruskal-Wallis test. Spearman correlation coefficient was used to evaluate the relationship between lecture capture analytics, VLE hours and individual attainment data. However, the paper does not present any individual student data points in case these points can be identified by the students. Therefore, while individual student data points were used to evaluate the correlation between lecture capture analytics, VLE hours and academic attainment, these data are presented by ranking the cohort based on the specific analytics parameter and then dividing these into 4 groups of approximately equal size (the same analytics value was not split across the banding). When students were divided into high and low engagement groups based on when they had accessed their lecture captures, their examination grades were first normalised by taking log_10_ of each data point, before testing whether there was a significant difference using Students’ unpaired t test. For analysis of the interrelationship between total lecture capture views, VLE hours and academic attainment the Kruskal-Wallis test was used. For all statistical analysis significance was assumed at the 5% level. Note that for some data sets, the small number of students in some groups precluded statistical analysis. In these cases, the data were reviewed in terms of trends.

### Ethical approval

Ethical approval was obtained from the University of Sheffield. The project code was 023251. Given the use of data collected for a different purpose in the study, approval was given to take a consent approach where students were informed about the research via Google+ communities, with a number of reminder e-mails and announcements in teaching sessions. Students then had the option to opt out of the research via e-mail at any point until the data was anonymised. Given this approach, no individual student data is reported.

## Results

The results section is broken down into 3 sections.

1. Whole second year cohort use of lecture capture.
2. Data evaluating the relationship between unique and total views of lecture captures and academic attainment and an analysis of the pattern of lecture capture use.
3. Data evaluating the relationship between VLE hours, total lecture capture views and academic attainment.

### Section 1: cohort use of lecture captures

The cumulative number of views of captures increased over the academic year, with a total of 50,603 lecture capture views from all students by the end of the year, figure 1A. By week 4 of semester 1, only 17 (10%) of the cohort had not watched any lecture captures. This number decreased across the academic year; 15 students at week 6, 9 at week 9, 8 at week 12, 6 the day before the January exam, 1 student by week 4 of semester 2 and by week 6 of semester 2 all students had accessed at least one lecture capture.

**Figure 1:**
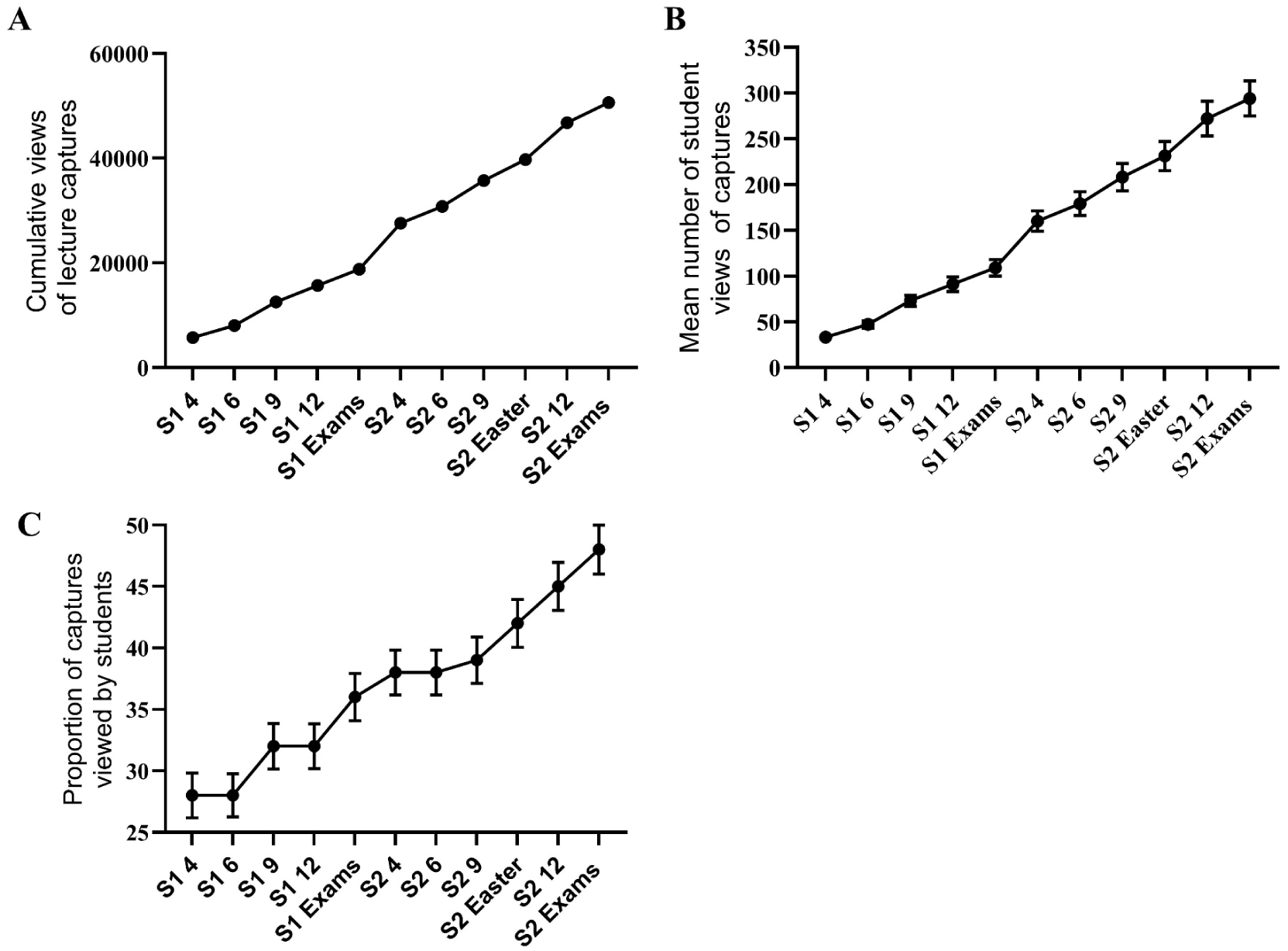
Mean cohort behaviour over the academic year. S1 = semester 1, S2 = semester 2. 4, 6, 9 and 12 refer to weeks of teaching (1 semester = 12 weeks). Exams = data collected just before the respective examinations. Easter = data collected at the end of the Easter vacation. A. Cumulative views of lecture captures across the academic year. B. Mean student lecture capture views. C. The proportion of unique lecture captures viewed by students.

As expected with additional lecture captures being available across the year, the mean number of total number of views for students increased across the academic year, figure 1B, P < 0.0001 (n = 172). There was no obvious increase in mean views just before each examination, suggesting that the cohort had not demonstrated massed learning behaviours, i.e. there was no evidence of cohort binge watching just before the examinations.

As students often viewed a lecture capture more than once, analysis was also done on the number of unique captures viewed across the academic year (expressed as a percentage of the total), figure 1C. Across the year, the proportion of captures viewed by the cohort increased, P < 0.0001 (n = 172). The proportion of captures viewed over semester 1 was 28 ± 2 % in week 4, versus 36 ± 2 % just before the exam (n = 172). There was a suggestion of possible binge watching behaviour between the end of semester 1 and just before the exam, with an uplift in the proportion viewed. In semester 2 up to week 9, the proportion was similar to the start of the semester, but there was then a trend for the proportion to increase from week 9 through to just before the exam, e.g. 38 ± 2 % versus 48 ± 2 % (n = 172) in S2 week 4 versus S2 exams, respectively.

### Section 2: lecture capture engagement and academic attainment

#### Is there a correlation between the total number of lecture capture views and academic attainment?

There was a significant, positive correlation between the total number of views of lecture capture and attainment in the MCQ examination for all time points in semester 1 (n = 168, P ≤ 0.007 for all), figure 2A. There was also a significant, positive correlation for all time points across both semesters for the SAQ / essay examination (n = 165, P < 0.04 for all), figure 2B. However, the total number of lecture capture views was not correlated with the coursework grade for all time points across the year (n = 169, P > 0.99, data not shown).

**Figure 2:**
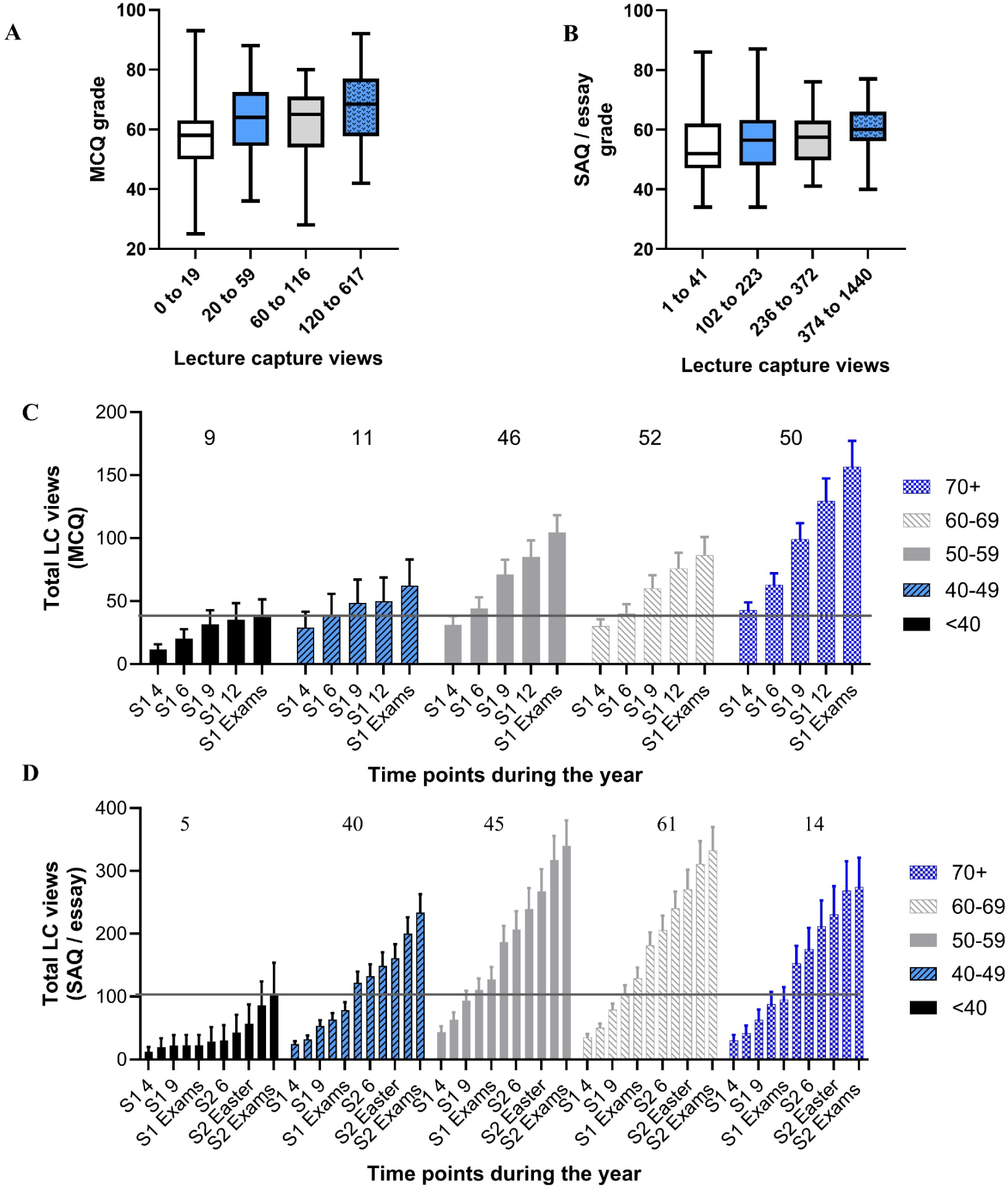
Relationship between total lecture capture views and academic attainment. A. MCQ grade versus lecture capture views in semester 1 week 12 (n=168). B: SAQ / essay grade versus lecture capture views in semester 2 week 12 (n = 165). Students were ranked based on views and subdivided into 4 groups. C: Mean lecture capture views across semester 1 for students in MCQ attainment bands. D. Mean lecture capture views across the academic year for students in SAQ / essay attainment bands. The horizontal lines in C and D show the maximum mean value for the fail students. Student numbers in each group are shown above each grade band.

When students were placed into MCQ attainment groups, there was a trend for the lower achieving students to have viewed fewer captures initially in semester 1, figure 2C, and show a smaller increase over the semester compared to the higher achieving students. Students achieving 50 or higher appeared to show a gradual increase in the number of views across the semester compared to those students who achieved 49 or lower.

When students were placed into SAQ / essay attainment groups, there was also a trend for the lower achieving students to have viewed fewer captures across the whole of the academic year, figure 2D. Students with grades below 40 seemed to show a relatively constant number of views across the academic year until just after the Easter vacation, and then there was a marked increase in views up to their assessment. Those students with 40-49 started with a number of views that was perhaps slightly smaller than the higher achieving cohorts. They had a gradual increase in views over semester 1, appearing to just lag behind the number of views for those students achieving 50+ in the SAQ / essay examination for all time points. They demonstrated a large increase in views by week 4 of semester 2, but this increase did not bring them up to the number of views of the students attaining 50+ in the examination. There was an additional increase in views in week 12 of semester 2, and just before the examination. While the 40 to 49 students finished the academic year with more views than those students who achieved below 40, they had fewer views than those students achieving 50+. There were no obvious differences between the 50-59, 60-69 and 70+ data, other than those students attaining 70+ may have had slightly fewer views compared to those attaining 50 to 69, and they appeared to have completed the majority of their views by the end of the semester.

#### Is there a link between the number of unique lecture captures that have been viewed and academic attainment?

There was a significant, positive correlation between the number of unique lecture captures viewed by students and attainment in the MCQ examination at all points in semester 1 (n = 168, P < 0.005 for all), figure 3A. There was also a significant, positive correlation for all time points across both semesters for the SAQ / essay examination (n = 165, P < 0.04 for all), figure 3B. However, the number of unique captures viewed by students was not correlated with their coursework grade (n = 169, P > 0.50, data not shown).

**Figure 3:**
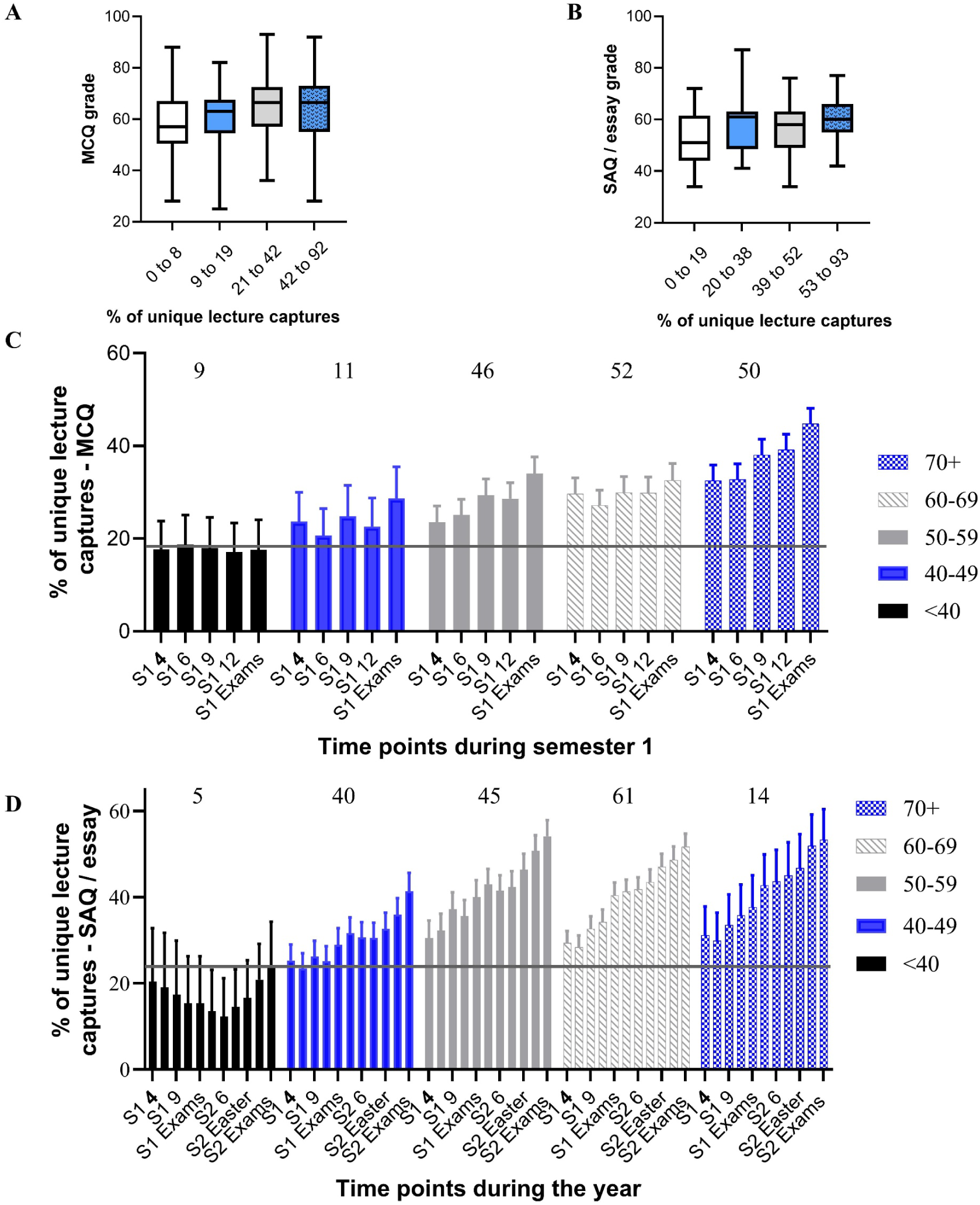
Relationship between unique capture views and academic attainment. A: Unique views in week 6 of semester 1 and MCQ performance. B: Unique views in week 6 of semester 2 and SAQ / essay performance. Data for A and B were banded into 4 groups based on number of lecture captures views (% of the total available). C: Mean unique views across semester 1 for students in MCQ attainment bands. D. Mean unique views across the academic year for students in SAQ / essay attainment bands. The horizontal lines in C and D show the maximum mean value for the fail students. Student numbers are shown above each grade band.

Lower achieving students in the MCQ assessment showed a trend for having viewed fewer unique captures over semester 1, figure 3C. They had typically viewed only 17.7 ± 6.1 % of the available captures in semester 1, and this did not change across the semester (17.6 ± 6.5 %). The highest achieving students on the other hand, had viewed 32.5 ± 3.4 % of captures early in the semester, and this rose steadily over the semester to 44.8 ± 3.4 % just before the examination.

There was also a trend for the lower achieving students in SAQ / essay assessment to have viewed fewer of the captures across the academic year, figure 3D. Students with fail grades seemed to show a drop in the proportion of unique captures viewed over semester 1 and early semester 2 (i.e. captures were made available but were not being viewed), but then a large increase in the run up to their semester 2 examination. Students with 40-49 had a higher number of initial views, but over semester 1 the proportion of unique captures being watched didn’t really change (25.3 ± 3.7 % versus 29.0 ± 3.9 %, in week 4 of semester 1 and just before the semester 1 exam period, respectively). They did show an increase at week 4 of semester 2 (31.7 ± 3.7 %), but again there was minimal further increase up to the end of the Easter vacation, 33.0 ± 3.7 %. From the end of the Easter vacation the number of unique views increased to 41.4 ± 4.2 % just before the SAQ / essay examination. There were no obvious differences between the 50-59, 60-69 and 70+ student data. These students viewed a higher number of unique captures and demonstrated a steady increase in the number viewed over the academic year. The 70+ students appeared to have completed their lecture capture views by the end of the semester.

Another way of reviewing the data is to group students within their attainment bands based on the number of unique views in relation to the cohort median, i.e. 1. Zero lecture captures viewed; 2. Lecture capture views were less than the cohort median and 3. Lecture captures views equal to or higher than the cohort median. These were then presented as a heat map (percentages of students in each group), figure 4. For the MCQ, higher achieving bands tended to have more students in the group that had unique views equal to or above the cohort median. The lower attainment bands tended to have more students in zero views or views below the cohort median. For the SAQ / essay heat map a similar profile was observed. In the 50 to 69 bands more students had unique views that were equal or higher than the cohort median. At 70+ there was a 50:50 split across equal / above and below. Below 49 more students were in the below cohort median grouping.

**Figure 4:**
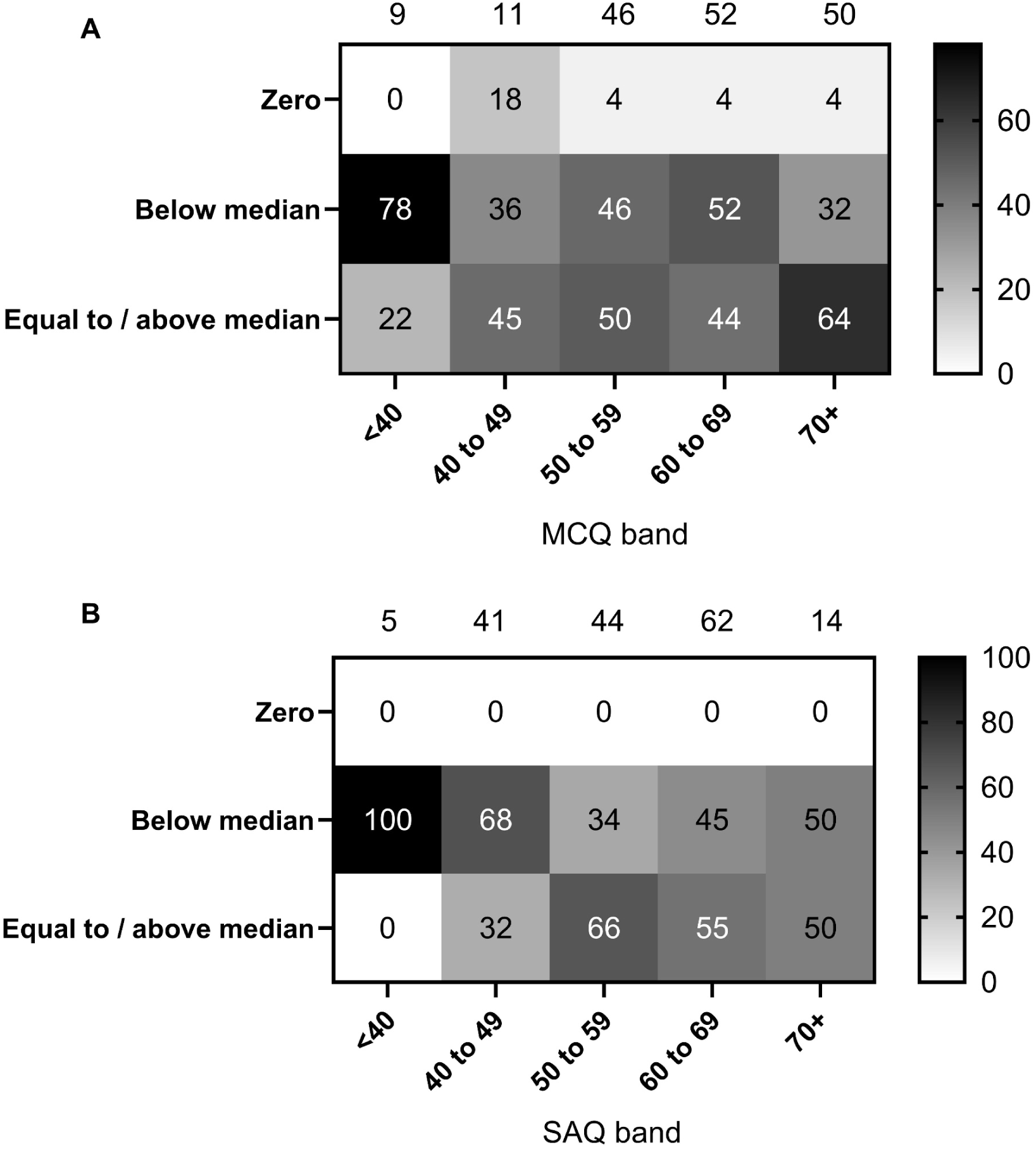
Heat map of unique lecture capture views analytics for week 12 for the MCQ (semester 1) (A) and SAQ (semester 2) (B) assessments. Numbers in each box are the percentage of students within the attainment band in each of the three groups. The numbers above each figure are the total number of students.

#### Is there a link between the pattern of engagement of lecture capture views and academic attainment?

The following section explored the pattern of engagement with lecture capture, irrespective of the total number of views.

The total views of a student in week 12 of semester 1 (MCQ) or semester 2 (SAQ / essay) was expressed as a percentage of their individual total views just before the relevant examination. Students were then placed into two groups, and the examination grades reviewed. 1. Students who had watched 70% or more of their individual views by the end of the semester (higher engagement). 2. Students who had either no views or had watched less than 70% of their individual views by the end of the semester (lower engagement). This was repeated for 60% and 50% views completed by the end of the semester. The hypothesis was that students with a lower engagement during the teaching period would show a lower academic attainment than those students who had spaced their use of lecture captures out across the teaching weeks.

For the MCQ examination, there was no significant difference between the mean grade attained across the 2 groups for all percentages, figure 5A (P = 0.66 to 0.37, n = 168). In contrast, for the SAQ / essay examination students who had viewed at least 70%, 60% or 50% of their individual total lecture capture views by the end of semester 2 achieved higher grades than those who had not, figure 5B (P = 0.02 to 0.03, n = 165).

**Figure 5:**
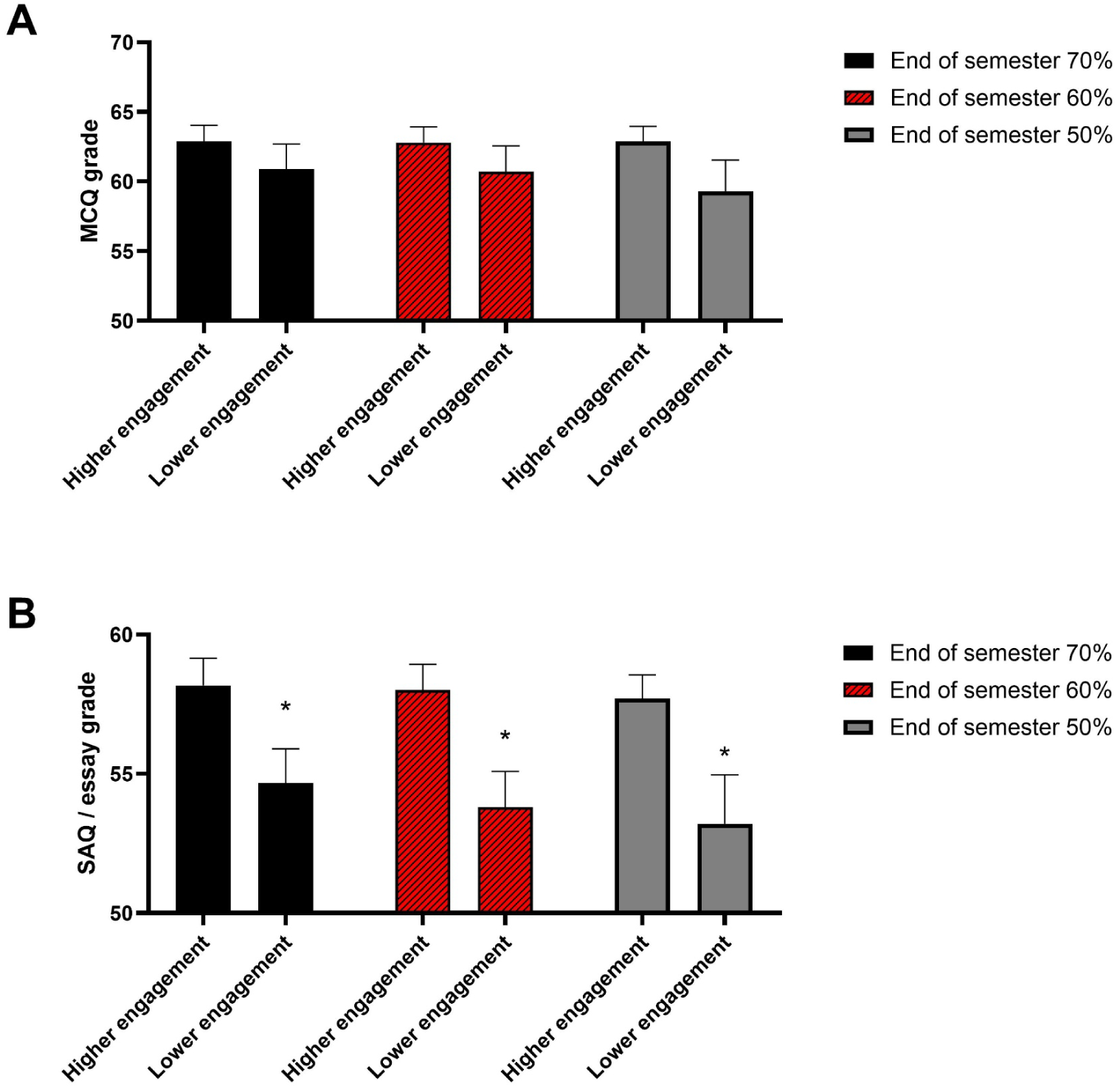
Relationship between the pattern of lecture capture views and academic attainment. A. Data for the year 2 MCQ examination. B. Data for the year 2 SAQ / essay examination. * indicates a significant difference.

For the SAQ / essay examination, just after the Easter vacation, there was also a significant difference in attainment between students who had completed 50% or more of their lecture capture views, 57.9 ± 0.9 (n = 122) versus 53.7 ± 1.33 (n = 43), for high and low engagement groups, respectively, P = 0.018. However, there was no significant difference for those with 70% or 60% views completed.

#### Is there a link between VLE hours, total lecture capture views and academic attainment?

Hours spent on the VLE was positively correlated with academic attainment for the MCQ (n = 168, P < 0.0003) and SAQ / essay (n = 165, P < 0.002) assessments, for all time points, figure 6A and B. It was also positively correlated with total lecture capture views, P < 0.0002 for all time points up to and including semester 2, week 12. For the day before the exam P = 0.01 (n = 172, data not shown).

**Figure 6:**
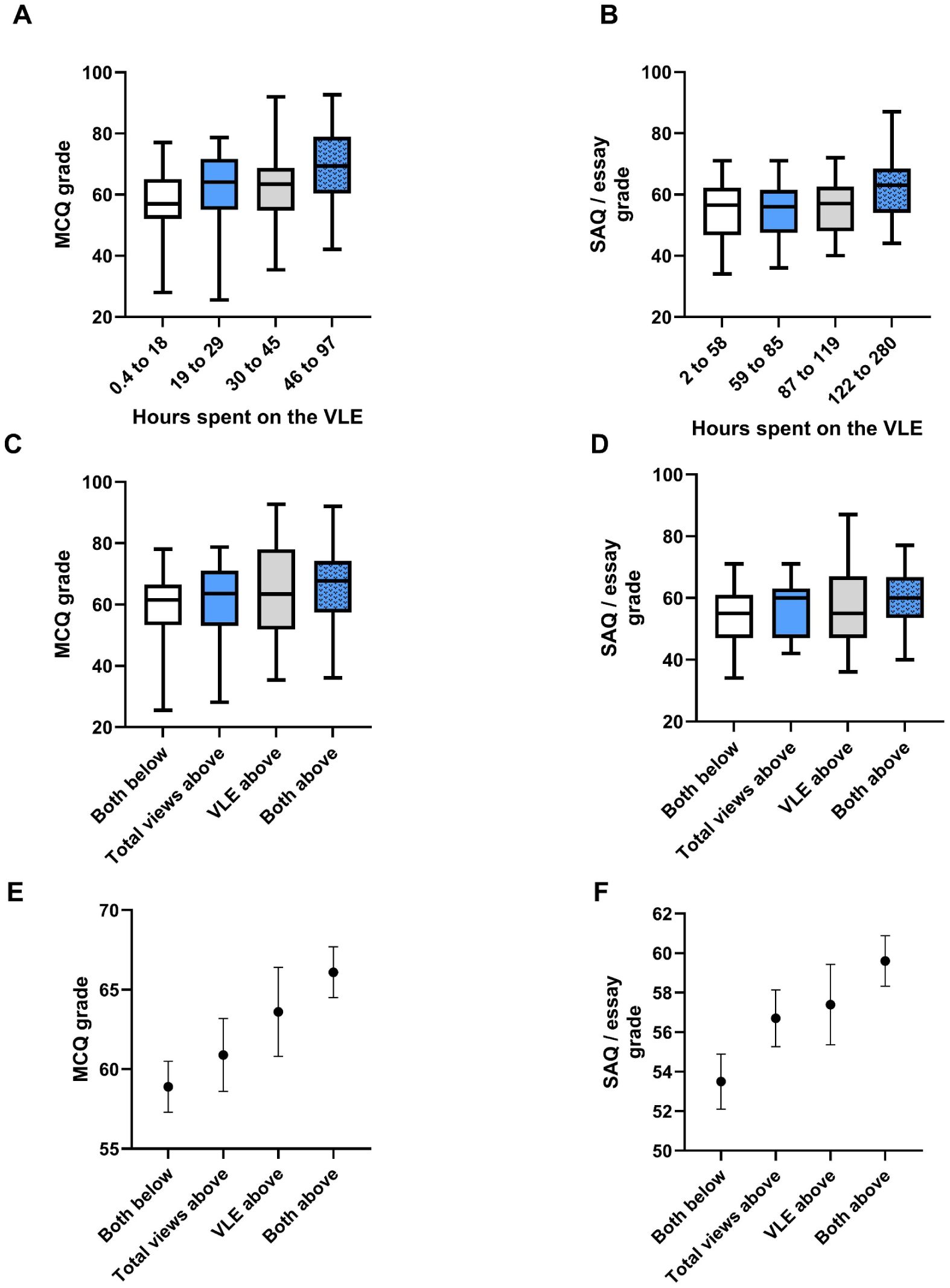
Relationship between VLE hours, total lecture capture views and academic attainment. A. MCQ grade versus VLE hours in semester 1, week 6 (n=168). B: SAQ / essay grade versus VLE hours in semester 2, week 6 (n = 165). For A and B students were ranked based on VLE hours and subdivided into 4 groups. C. (MCQ) and D (SAQ / essay) grades for students placed into groups based on whether they were below or above the cohort median. E. (MCQ) and F. (SAQ / essay) mean grades for students in these groups.

To evaluate the interrelationship between VLE hours, total views and academic attainment, students were placed into four groups based on whether VLE hours and total lecture capture views were equal to / above or below the cohort median. The attainment for each group was then compared. The specific groups were:

1. Both parameters below the cohort median.
2. VLE hours below, lecture capture views above the cohort median.
3. VLE hours above, lecture capture views below the cohort median.
4. Both parameters above the cohort median.

There was a significant difference in the MCQ attainment of the four groups, Kruskal-Wallis, P = 0.02, figure 6C. The attainment for students below the cohort median for both parameters (n = 56) was significantly lower than the attainment for the group who were above the cohort median for both parameters (n = 56), P = 0.007. While statistical analysis did not show any significant difference across the other groups, was a trend for the top and the lowest grade in the groups with above median for VLE hours or both parameters to be higher than those in the other groups. There was also a trend for mean grades to be ranked low to high in the following order, both parameters below median (n = 56), total views above median (n = 28), VLE hours above median (n = 28) and both above median (n = 56), figure 6E.

There was also a significant difference in the SAQ / essay attainment of the four groups, Kruskal-Wallis, P = 0.03, figure 6D. The attainment for students below the cohort median for both parameters (n = 47) was significantly lower than the attainment for the group who were above the cohort median for both parameters (n = 48), P = 0.01. Again, there were some interesting observations, although these were less clear cut than the MCQ data. There was a trend for the top grade in the groups with above median for VLE views or both parameters to be higher than the other groups (this was more obvious for the VLE above median cohort). In contrast to the MCQ data, the lowest grade in each group did not show any pattern. However, there was a trend for the mean grades to be ranked low to high in the following order, both parameters below median (n = 47), total views above median (n = 35), VLE hours above median (n = 35) and both above median (n = 48), figure 6F.

## Discussion

This study has evaluated the value of lecture capture use as a marker of engagement for student success. Overall, the findings from the study suggest that student use of lecture captures can be a potential engagement marker for student success in examination assessments, although it cannot be used in isolation, and should be considered in the context of other engagement markers. However, there are some important nuances in the data that must be considered and these will be discussed below. The study also reinforces that engagement with the VLE is associated with higher academic attainment, as shown in a number of other studies [35–38]. The study also demonstrates that engagement with both the VLE and lecture capture is correlated with the highest performance, i.e. there are benefits for students to engage with more than one learning resource.

Before considering the data in detail, an important point to make is that these data provide no evidence that the use of lecture capture has directly improved grades. The data simply suggest that student engagement with lecture capture is positively correlated with attainment, i.e. this is about engagement with learning. One of the students who completed a previous ethically approved survey that asked students about their perception of lecture capture (unpublished data) put this very nicely

> P1: “Are academically successful and motivated students more likely to engage in all revision sources, so when they attain high grades give the illusion of having being helped by lecture capture, when in fact they were going to achieve strongly regardless of their method of revision?”

While a number of previous studies have tried to evaluate whether lecture capture impacts on attainment, there is no consistent finding [25–30]. The impact of lecture capture on students is complex, and integrated with many other factors, including whether they have additional support needs, other personal challenges, how they use the captures, how they engage with their studies using other resources, their previous lived experience of learning. For some students, access to captures is likely to be positive, for others it may have a negative impact if they are not using the captures effectively. Therefore, there is no “one size fits” all when it comes to the impact of lecture capture, and we should therefore anticipate variation in the findings reported on the impact of lecture capture on student learning.

Having clarified that these data are looking at lecture capture use as an engagement marker, what do these data actually show us? The most obvious finding is that at a cohort level, for both examination assessments, the total lecture capture views and number of unique captures viewed were positively correlated with attainment. Students who did poorly in their examinations were less likely to have watched the lecture captures during the teaching period, i.e. they didn’t space their learning. For those students who failed the assessment the mean number of total and unique views were the lowest out of all the attainment groups. These students also demonstrated evidence of massed learning, with large increases in views just before the SAQ / essay exam in particular. Spacing learning as part of a self-regulated learning approach is known to be associated with academic attainment [40, 41]. Therefore, it is perhaps not surprising that the students in this study who demonstrated a massed or binge learning behaviour struggled in their assessments. Those students attaining grades of 40 to 49 had more use of lecture captures, but still below those students in higher attainment groups. Again, these students were more likely to have use that sat below the cohort median, although this was less obvious for the MCQ assessment. They also demonstrated massed behaviour, with the largest increase in lecture capture use typically in the run up to the examinations. These two cohort data sets therefore show lower use of lecture captures overall, i.e. they are consistent with lower engagement.

For the remaining attainment groups, the use of lecture captures was similar, with just a few small differences, suggesting that for these students it is more challenging to identify how lecture capture as an engagement marker indicates which students are likely to achieve the highest grades. For total and unique lecture capture views, those in the highest attainment group for the MCQ assessment had the largest total number of views, while the 70+ cohort for the SAQ / essay assessment seemed to have fewer total views than those attaining 50 to 69. The reason for this is not clear, but may reflect the differing nature of the assessments, that the highest achieving students used the captures more effectively to support their learning, or that those that engaged more with the semester 1 lecture captures did not need to revisit this content as much during the rest of the academic year. Indeed, 8 out of the 14 students attaining 70+ in the SAQ / essay assessment had also attained 70+ in the MCQ assessment, i.e. were part of the cohort of 50 students with the largest lecture capture use in semester 1. The other nuanced aspect of the data is related to the change in lecture capture use between the end of the semester 2 teaching period and just before the assessment. For the cohort attaining 70+ there was only an increase of 5.5 in the mean number of total lecture capture views, which was much smaller than the other cohorts over the same time period. This is consistent with these students having completed the bulk of their studies by the end of the teaching period, leaving the revision time for consolidation of learning.

How do these data compare to previous studies? Nordmann et. al. (2019) evaluated the relationship between the number of lecture capture minutes watched and exam performance for Psychology students [29]. A positive correlation was observed between minutes and exam performance for years 1 and 2, while for year 3 and 4 no correlation was observed. While the early year data is similar to the current study, the students that are at the same level in the Nordmann study are actually the year 3 students, where no correlation was observed. The data in the current study are therefore different, although the parameters evaluated were total views and unique number of captures watched rather than total minutes. The current study also mapped across the academic year, rather than a single time point. Edwards and Clinton (2019) also looked at a year 2 cohort, observing no correlation between lecture capture views and exam performance, although they did see a positive correlation between term time viewing and coursework, which is different to the current study [27]. A lack of correlation was also observed when lecture capture views at the end of the assessment period was compared to exam performance in year 2 and 3 biology students, [33]. This study did not evaluate use during the teaching semester, and therefore could not differentiate between students who had binge watched and those who had spaced their use across the teaching semester. In contrast, Williams et. al. (2016) saw a slightly positive correlation between lecture capture views and predicted exam performance [32], although this was in a first year biology cohort (equivalent to the year 2 cohort in the Nordmann et. al. (2019) study where a positive correlation was also observed, [29]).

In at least one paper, the difference to the current study might be explained by how the students in those studies used lecture captures [27]. In that study, where no correlation to the exams was observed, 2/3rds of the use was in the revision period after teaching had been completed. This contrasts with the current study, where 84% and 88% of views were completed during semester 1 and semester 2 teaching, respectively. Therefore, students in the current study had spaced their use of lecture captures across the teaching semester, while in the Edwards and Clinton study students took a massed learning approach. In the Williams et. al (2016) study, where a small correlation was observed to exam performance, 44.6% of all views were completed in the 48 hours before an exam, suggesting a massed learning approach by some students [32]. They did however observe higher than predicted exam attainment in students who had accessed the lecture captures early in the teaching period, while those who accessed the captures later tended to do less well than predicted. These data align with the current study, which shows that early engagement was associated with better examination performance. A study in 2024 also identified that those who watched the captures during the teaching semester tended to do better in the January and final year assessments [34].

Our student cohorts are, of course, a diverse group, with individual strategies and experiences, which all impact on their learning behaviour and we know that not all students use lecture captures effectively. This variation within the different cohorts can be seen in the various box plots, e.g. looking at the maximum and minimum values, and also in the heat map data. In the box plots, if we look at those with the lowest number of total lecture capture views, their grade in the MCQ and SAQ / essay assessments ranged from 93 to 25 and 86 to 34, respectively. Therefore, there are students in this group who attained high grades in the assessments with low engagement with lecture capture views. This finding is not surprising. There will always be high achieving students who have good learning strategies that do not require them to have a high engagement with lecture capture. Looking at the SAQ / essay data reveals that of the 41 students with the lowest total number of views of lecture captures only 3 attained 70+ (7%), while 17 (41%) attained 49 or lower. If we look at the SAQ / essay heat map we can see the trend for more students with lower grades to be below the whole cohort median, but what is also clear is that even in the highest attainment band there are students that sit below the median. At the two extremes, those who failed this assessment all sat below the median, while in the 70+ group there was a 50:50 split. This means that there are students who are achieving high grades in their assessment, but with low use of lecture captures. If we look at the relationship between VLE hours and total views we can see that 6 of the 14 students who attained 70+ in the SAQ / essay assessment were above median in both parameters, 4 above for the VLE hours and 2 above for lecture capture views (data not shown). Interestingly, 2 of the students were below the median for both parameters. Similarly, there are students with lots of views but worse academic attainment. For these students it is likely that they were not using the lectures effectively as part of their learning strategy (although there is no evidence that can be presented to support this statement). Some students in the lower attainment groups with larger views showed a large increase in views in the last few weeks of the semester and into the exam period, i.e. they took a massed learning approach, which may have contributed to their relative lack of performance. There were 45 students attaining 49 or lower in the SAQ / essay assessment. Looking at their VLE hours and total views, 8 were above the median for both parameters, 11 were above for VLE hours, 10 above for lecture capture views and 16 below for both parameters. Therefore, some of these students are using the VLE resources and lecture captures, but are potentially less aware of what best practice in learning looks like.

These data are consistent with the whole cohort analysis and discussion, but do highlight the importance of taking each student as an individual when evaluating engagement with lecture capture resources, i.e. there are outliers in all groups, and we cannot simply take a single measure of engagement if we want to track students for early support and intervention.

The final area for discussion is related to the timing of when a student watches the lecture captures, irrespective of the total number of views a student has. For example, two students might have a total of 10 and 100 views each. If they had completed 70% of these by the end of the semester (7 and 70 views, respectively) they were placed into the same engagement group. The idea behind this approach was to normalise views to the total for an individual, rather than the whole cohort, and then map when these views were completed. It was clear that, for the SAQ / essay examination, higher grades were obtained by those students who had spaced their use of the captures out, irrespective of the total number. The data showed that at the end of the semester, higher grades were obtained by those students who had completed 50%, 60% or 70% of their views by this time point. Indeed, students who had completed 50% of their total lecture capture views by the end of the Easter vacation attained higher grades than those who had not. These data suggest that students with lower number of lecture capture views, but who space their views out across the teaching period are also more likely to do well in assessments. Interestingly, the same relationship was not present when considering the MCQ assessment. There was no significant difference between the lower and higher engagement groups. The reason for this difference is not clear, but may be related to the fact that the SAQ / essay assessment spanned the whole academic year, while the MCQ assessment was only semester 1. It is likely that trying to take a massed learning approach for whole year content (SAQ / essay) was more challenging than doing this for a single semester (MCQ).

Overall, these data are consistent with the use of lecture captures providing a potential marker for student engagement. The work adds to the body of literature in this area, and is the first to directly evaluate the relationship between engagement with lecture captures and academic attainment at time points across a whole academic year. The study shows that in terms of the lowest performing cohort, there is a clear link between low engagement with captures during the teaching period, along with a massed approach closer to the examinations. While the use of lecture capture as an engagement marker to identify the higher achieving students is less clear, there are still valuable nuanced aspects in the data, that can be used to highlight to students the importance of spacing their use of lecture captures out across the teaching period. The study shows that lecture capture analytics have the potential to help academic staff identify those students who may be struggling with their studies, in particular if the analytics are combined with other engagement markers such as the use of the VLE.

## Acknowledgements

The author declares the use of Notebook LM to complete an initial summary of published papers identified from Google Scholar. These summaries were used to identify the key papers to read and review, to use in the current paper.

## Disclosure of interest

The author reports that there are no competing interests to declare.

## Notes

### Competing Interest Statement

The authors have declared no competing interest.

